# Auditory-motor synchronization determines the use of predictions in music perception

**DOI:** 10.1101/2025.08.20.671291

**Authors:** Nina Hartung, Alice Vivien Barchet, Cornelius Abel, Benjamin R. Pittman-Polletta, Fredrik Ullén, Johanna M. Rimmele

## Abstract

When listening to music, our brain constantly generates predictions about the timing and pitch of upcoming sounds based on the structure of the music. Individual differences in the ability to make such rhythmic and melodic predictions shape music perception. Previous studies show that during rhythmic auditory-motor synchronization tasks, better performance is related to stronger motor system engagement. However, whether individual differences in auditory-motor interactions influence higher-level rhythmic predictions is unknown. Here, we used electroencephalography during a naturalistic music listening task to assess the neural tracking of rhythmic and melodic predictions generated from (short-term) local contextual information and (long-term) experience using the Information Dynamics of Music model. We combined a multivariate temporal response function approach with two behavioral measures of auditory-motor synchronization (whispering and finger tapping). We found that stronger auditory-motor synchronization predicted stronger neural tracking of lower-level temporal aspects of the music signal, i.e. acoustic envelope and note onset. Across participants, higher-level (short- and long-term) predictions were also neurally tracked, with neural tracking stronger for rhythmic than melodic predictions. Importantly, our evidence suggests individual differences in the weighting of musical predictions. Individuals with stronger auditory-motor synchronization showed stronger neural tracking of rhythmic compared to melodic predictions, and this effect was especially pronounced for short-term predictions. Our findings demonstrate that the motor system is critically involved at several processing levels even in purely perceptual music tasks, and pave the way for understanding individual differences in the weighting of rhythmic and melodic predictions during music listening.

**Significance statement:** Our ability to make musical predictions affects our perception of music. Understanding individual differences in predictive processing may allow for differential diagnostics and interventions in clinical disorders such as rhythm disorders or hearing impairment. In an electrophysiological study, we found evidence that the motor system is implicated in predictive processes during perception of music. Specifically, we find advanced neural processing of acoustic temporal features and stronger weighting of short- and long-term rhythmic musical predictions in individuals who recruit the motor system more strongly. Likely, in individuals with high compared to low auditory-motor synchronization strength, the motor system facilitates temporal processing at different levels: at the level of acoustic temporal structure processing, and at higher-level processing of rhythmic musical predictions.

## Introduction

Music is universal across cultures (1), and humans show a natural predisposition to perform and appreciate it (2, 3). Given its role in human social interactions, musical engagement has been suggested to have beneficial effects on our wellbeing (4). The currently prevalent idea that perception is essentially shaped by predictive processes (5) has been extended to music (6). As we listen to a musical piece unfolding over time, predicting the upcoming notes, harmonies, and rhythm is central for an adequate perception of the structure of the music (6–12). Indeed, our tendency to move to music and enjoy it is related to the extent to which we can predict musical structure (11, 13).

One approach to analyzing the neural processing of rhythm has been to characterize neural tracking. This essentially refers to quantifying the alignment of neural activity with, for example, the acoustic envelope, and reflects slow rhythmic modulations at the single note and beat level (14–16). Envelope tracking has been related to temporal segmentation based on neural oscillations (15). Although this may allow for the processing of quasi-isochronous rhythms, it cannot directly account for more complex predictions based on musical structures (17). Predictions of multiple features such as the temporal onsets and pitch of notes have been shown to be formed in parallel (18–20). An important distinction in this context has been made between *short-term prediction*, which is based on statistical learning within a musical piece (21, 22), and prediction based on long-term experience (*long-term prediction)*. The latter takes statistical features of a whole corpus of music into account, and thus is based on enculturation and life-long experience with particular types of music (8, 19). Recent data support that humans can extract both short-term and long-term predictions during music listening (18). For example, listeners have been shown to extract both rhythmic and melodic predictions for Western classical music (23–27). A recent study showed that individuals are extracting melodic predictions based on implicit long-term knowledge (priors) but, at the same time, are sensitive to short-term regularities within the current musical piece – as reflected by neural tracking (23).

Musical predictions can be flexibly top-down modulated – that is, the commitment to a particular prediction can be up-weighted or down-weighted by adjusting its strength – depending on the musical context (precision weighting, 13, 18–20). As a comparison, in speech, the utilization of semantic predictions has been suggested as a compensatory strategy to facilitate comprehension in acoustically demanding listening situations (30, 31). How different types of predictions are integrated during music perception, however, remains unclear. A recent study (26) found no difference between the neural tracking strength of melodic and rhythmic predictions during passive music listening, suggesting that both were utilized. However, individual differences in the weighting of predictions were not investigated. Cantisani et al. (32) studied the neural tracking of pitch and semantic predictions and reported trade-off effects when listening to songs, in favor of pitch predictions over semantic predictions. The appearance of these trade-offs may have depended on both song characteristics and participants’ preference. These findings suggest individuals may differ in the extent to which they are able to extract or utilize musical predictions. For example, individuals may differ in their specific prediction weightings, so that some people mainly rely on rhythmic predictions while others have a stronger focus on melodic structures, with the latter selectively up-weighting melodic predictions. Recently, individual differences in rhythmic temporal processing of speech (33, 34) and music (35–37) have been demonstrated. Moreover, advantages in temporal processing such as improved rate discrimination, duration deviant discrimination, and syllable recognition have been related to a stronger spontaneous recruitment of the motor system (34, 36, 38). Based on this (and other; e.g. 40) research, we expect increased motor system recruitment to enhance rhythmic processing at a lower-level such as the acoustic envelope tracking. Given the central role of the motor system (premotor cortices, supplementary motor area and basal ganglia) in musical rhythm and beat processing (3, 13, 40–45), one could hypothesize that individual differences in auditory-motor interactions might also affect the ability to extract rhythmic predictions. Indeed, the claim that the recruitment of the motor system is related to predictive processing is central to influential oscillatory and predictive coding theories (13, 46–48), and implicit in various studies. The link between motor system recruitment and the ability to make rhythmic predictions, particularly long-term predictions, however, has not been directly investigated. One the one hand, a crucial role of the motor system in rhythmic enculturation -the build-up of rhythmic prediction based on long-term experience-has been proposed (51). On the other hand, during rhythmic development the motor system may only gradually come into play during the first decade of life (52, 53), possibly resulting in a lesser impact of the motor system on long-term compared to short-term predictions. In addition to auditory-motor synchronization, musical training may be another potential source of individual differences in prediction weighting (32). Since musical training and auditory-motor synchronization strength are associated (35), both may possibly aid rhythmic predictions. Yet, an up-weighting of pitch predictions in musically trained individuals, as shown by Cantisani et al. (32), is also possible. However, despite growing interest in the topic, empirical studies of prediction weighting and its interindividual differences remain limited.

Based on these earlier findings, we hypothesize that stronger auditory motor-synchronization is related to enhanced neural tracking of temporal aspects of the music (*lower-level rhythm processing*; i.e. the acoustic envelope and note onsets), as well as an up-weighting of rhythmic over melodic predictions. To quantify neural tracking of predictions during naturalistic music listening (Fig. 1A), we utilized a multivariate temporal response function (mTRF (62–64)) approach (Fig. 1 B). Musical predictions were estimated with the Information Dynamics of Music model (IDyOM, (65, 66)). In our analyses, we included IDyOM models based on, firstly, local context only (“short-term”), and secondly, pre-trained long-term regularities (“long-term”). Individual auditory-motor synchronization strength was assessed in a speech-production-perception (“whispering”) synchronization task (33, 67) and an analogously designed finger tapping task (36). Self-report data on musical sophistication (68) was collected to allow us to control for the influence of musical training on long-term prediction weighting and its interplay with auditory-motor synchronization abilities. Additionally, the participants’ auditory working memory capacity was estimated as it may affect the ability to use long-term predictions (31) and the processing of music in general (69). Lastly, ratings of familiarity and enjoyment for each musical piece were included as control variables (25, 70, 71).

**Fig. 1:**
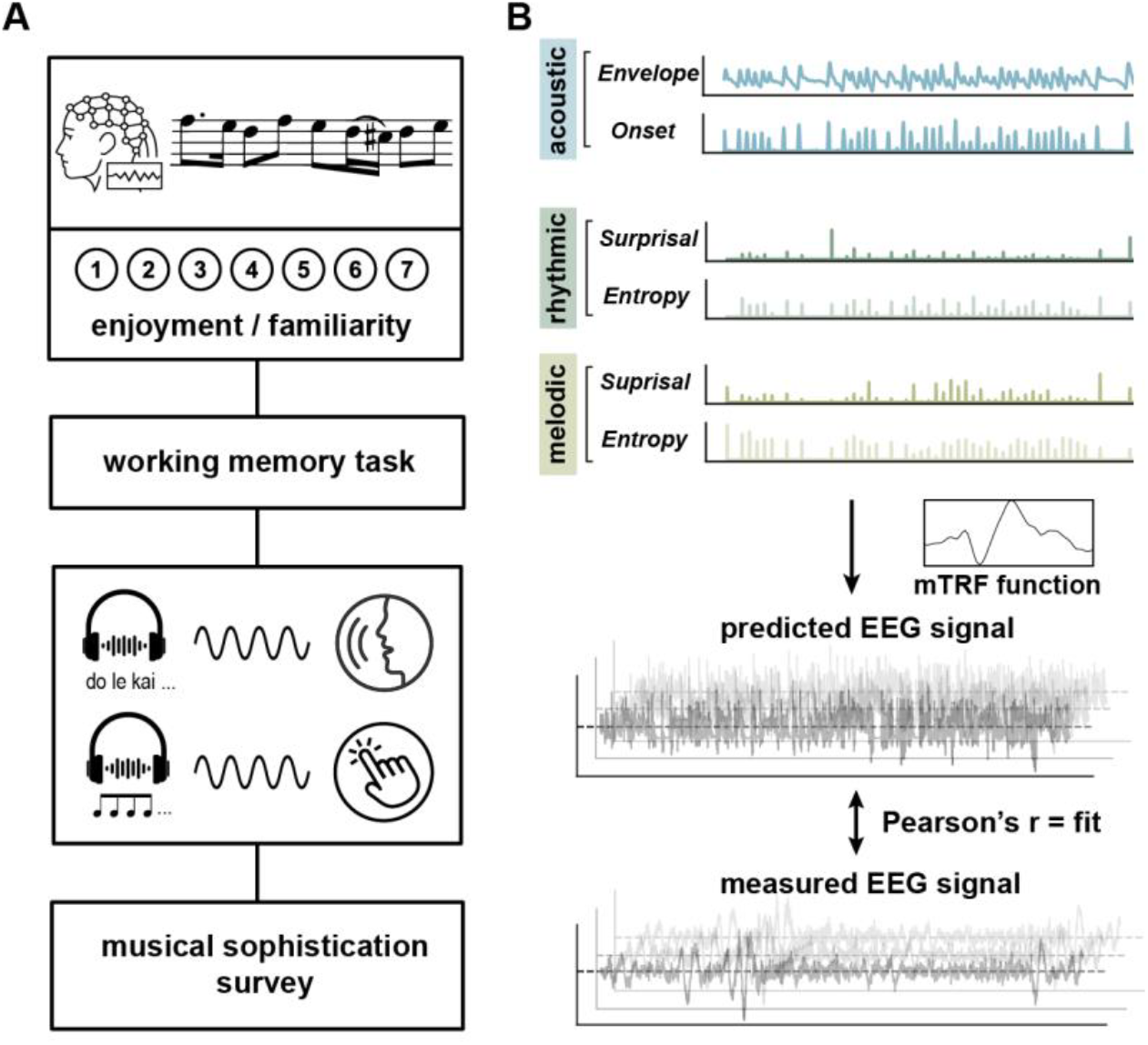
Overview of research paradigm. (***A***) Experimental procedure: Participants listened to 11 blocks of naturalistic, monophonic melodies (one melody per block) and entered an enjoyment as well as a familiarity rating after each melody. This was followed by an assessment of the auditory working memory span (Digit Span). Auditory-motor synchronization strength was measured using a whispering synchronization task and an adapted finger tapping task. Lastly, participants filled out a questionnaire on musical sophistication (Musical Sophistication Index, MSI). (***B***) Computational modeling of neural tracking: stimuli characteristics were quantified in a temporal acoustic feature-only model (“acoustic”) as well as two musical prediction models “short-term” and “long-term” both of which contained sub-models for melodic and rhythmic predictions. *Entropy* and *Surprisal* values quantified predictions and were estimated with IDyOM (see Methods for details). On their basis, neural tracking was computed using multi-variate temporal response functions (mTRFs) which are based on time-resolved linear regression. The resulting *mTRF fit* describes how well the mTRF predicted unseen EEG data based on the given stimulus features. The *mTRF fit* can thus be interpreted as measure for neural tracking of those features.

We found that individuals with stronger auditory-motor synchronization showed stronger neural tracking of temporal aspects of the music acoustics during naturalistic, passive music listening. Moreover, our results suggest that strong auditory-motor synchronization is likely to predict an up-weighting of rhythmic over melodic short- and long-term predictions on an individual level. Taken together, the recruitment of the motor system seems to be related to the facilitation of temporal processing of music, at both the level of acoustic temporal processing as well as higher-level rhythmic musical predictions.

## Results

Behavioral and neural data from 40 healthy participants were included in the analyses. From the mTRF approach, we retrieved measures for neural tracking quantified as *mTRF fit* of each model (Fig. 1B). Apart from the *mTRF fit*, each mTRF model constructs temporal response functions for each feature, also called weights or kernels, that describe the feature’s influence on the neural activity over time (i.e. the set time lag) (Fig. 1B). Generalized linear mixed-effects models (GLMM) were separately computed for the acoustic and prediction models to analyze the effects of auditory-motor synchronization (*PLV*) (with a GLMM for “whispering” and one for “tapping”), musical sophistication (*MSI*), working memory (*digit span*), and ratings (*enjoyment* and *familiarity*) on neural tracking (*mTRF fit*). Random intercepts for *subject, trial* and *EEG-electrode* were included. Using cluster-level statistical permutation tests (72, 73), electrodes with significant activity were identified and separately included in the GLMMs. (Note that our GLMM results do not change if a cluster is instead selected based on the previous literature, Supplementary tables S1-3 and S4-6.) For the sake of clarity, only the “whispering” GLMMs will be reported, given the higher variance in *PLV*s as opposed to the “tapping” *PLV*s. Note, the “tapping” GLMMs produced overall similar results and can be found in the Supplement (S7-9). Descriptive statistics of the control variables can be found in the Supplements.

### Tapping and whispering synchronization are correlated

As indicators of auditory-motor synchronization strength, phase-locking values (*PLV*s) between the acoustic and the produced signal were computed for the “whispering” synchronization (SSS test, n=36) and the “tapping” synchronization task (n=40). Performances were significantly correlated across tasks (n=36, Spearman’s rho: ρ=0.52, p=0.001). Visual inspection of the distribution suggests a characteristic bimodal shape (with high synchronizers, n=17, low synchronizers n=19; Fig. S10), as described previously (e.g. 20–22) for the *PLV*s from the whispering task. The *PLV*s of both tasks were included as continuous predictors. Due to multicollinearity between the *PLV*s, separate GLMMs were computed (“whispering” and “tapping” GLMMs).

### Auditory-motor synchronization positively predicts acoustic temporal feature tracking

Statistical cluster inference revealed a broad distribution of acoustic temporal feature tracking across the brain with significant differences from the (shuffled) baseline at all EEG-electrodes (Fig. 2A). Visual inspection suggests the strongest tracking at fronto-central sites, consistent with sources in the auditory cortex (e.g. 44–46). The mTRF functions for the “acoustic” model (acoustic envelope and note onset) depict peaks between 0 and 100 ms (Fig. 2A), indicating an early response to low level acoustic features as it has been reported previously (23, 77).

**Fig. 2:**
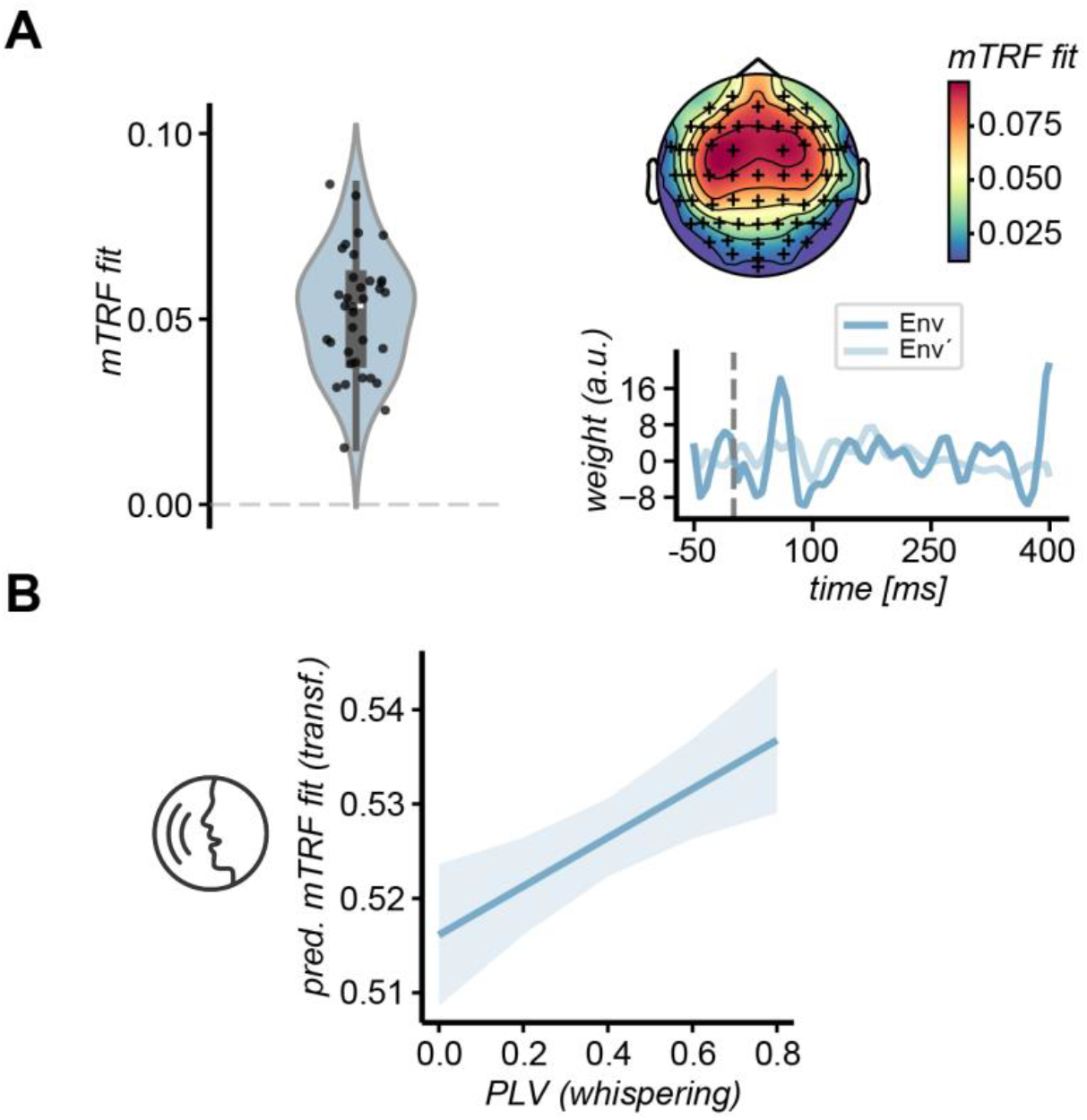
Significant neural tracking of acoustic features. Auditory-motor synchronization and enjoyment effects on acoustic feature tracking. (**A**) The figure displays the results of the mTRF analysis, including the *mTRF fits*’ distribution with topographies as well as the mTRF weight function for the “acoustic” model (“Env” = acoustic envelope; “Env’” = note onset). Box plots within violin plots indicate the median across subjects (white line), the quartiles (black boxes) and full distribution (whiskers). Electrodes included in a cluster with a significant mTRF fit are marked (+). (**B**) Statistical analysis revealing a positive effect of auditory-motor synchronization strength on neural tracking of acoustic temporal features. (siginificance levels *p* < 0.05)

A generalized linear mixed-effects model was used to predict the effect of auditory-motor synchronization, musical sophistication and their interaction on the neural tracking of the acoustic temporal features. The “whispering” GLMM revealed a significant fixed effect of the *PLV* (estimate=0.018, standard error=0.006, 95%CI = [0.007 – 0.029], pFDR=0.004). Hence, a higher auditory-motor synchronization predicted stronger neural tracking of the acoustic temporal features (acoustic envelope and note onset) (Fig. 2B). Additionally, a negative effect for *enjoyment* (estimate=-0.002, standard error=0.000, 95%CI=[-0.003 – -0.001], pFDR<0.001) was present, indicating higher neural tracking of acoustics when the *enjoyment* rating for the musical piece was relatively lower (S1). No effects of *MSI, digit span*, and *familiarity* were observed (see table S1 for detailed statistics). The “whispering” GLMM yielded a marginal R^2^ of 0.059 and a conditional R^2^ of 0.852.

### Auditory-motor synchronization determines the weighting of melodic and rhythmic predictions

*MTRF fits* tested against a (shuffled) baseline yielded significant neural tracking in fronto-central clusters in all mTRF models (Fig. 3A), likely reflecting sources in auditory cortex and frontal brain areas (e.g. 44– 46). In line with recent findings, the mTRF weights for all musical prediction mTRF models show peaks around 50 ms and 200 ms (Fig. 3A) which might reflect the early response to lower-level acoustic processing and higher-level processing of predictions (23), respectively.

**Fig. 3:**
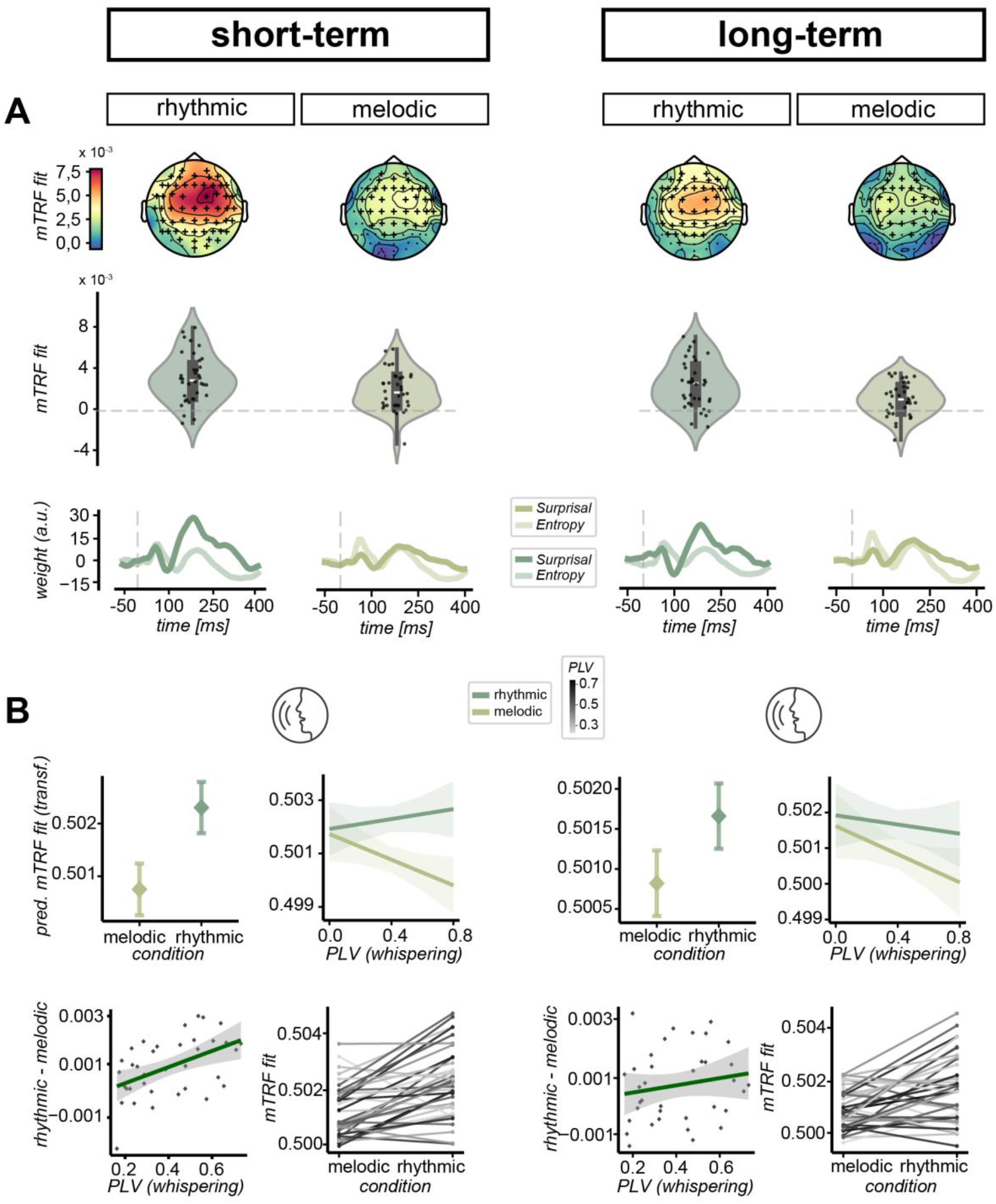
Significant neural tracking of short- and long-term musical predictions. A high PLV (“whispering”) predicts up-weighting of rhythmic over melodic predictions. (**A**) The figure displays the results of the mTRF analyses, including the *mTRF fits*’ distribution with topographies as well as the mTRF weight functions for the the musical prediction features. Box plots within violin plots indicate the median across subjects (white line), the quartiles (black boxes) and full distribution (whiskers). EEG-electrodes included in a cluster with a significant *mTRF fit* are marked (+). (**B**) GLMMs were used to predict the *mTRF fit*. In both, the “short-term” and “long-term” GLMMs, a significant main effect of *condition* was found, in that rhythmic predictions are neurally tracked more strongly than melodic predictions overall. A significant interaction shows that stronger neural tracking of rhythmic predictions is increasingly present with a rising *PLV* (whispering). In a post-hoc analysis, the contrast between rhythmic and melodic *mTRF fits* is significant in the “short-term” model. Additionally, subject-wise *mTRF fits* (of both *conditions*) are plotted, slopes are color-coded according to the individual *PLV*. (siginificance levels *p* < 0.05)

To investigate the weighting between musical melodic and rhythmic predictions, GLMMs were constructed that additionally included *condition* as fixed effect (coding for either melodic or rhythmic predictions). Hence, the neural tracking was predicted based on auditory-motor synchronization (*PLV*), musical sophistication (*MSI*) and *condition*, as well as their interactions.

The “whispering” GLMMs revealed a significant main effect of *condition* for both the “short-term” and “long-term” mTRF models (“short-term”: estimate=0.006, standard error=0.000, 95% CI=[0.006 – 0.007], p(FDR)=0.000; “long-term”: estimate=0.003, standard error=0.000, 95% CI=[0.003 – 0.004], p(FDR)=0.000), i.e. overall stronger neural tracking of temporal compared to pitch predictions (Fig. 3B).

Importantly, we found a significant interaction between the *PLV* and *condition* for both models (“short-term“: estimate=0.002, standard error=0.000, 95%CI=[0.002 – 0.003], pFDR<0.001) (“long-term“: estimate=0.001, standard error=0.000, 95%CI=[0.001 – 0.001], pFDR<0.001). This interaction suggests that a relatively high whispering synchronization relates to an up-weighting of rhythmic over pitch predictions (Fig. 3B). In a post-hoc analysis, we found a significant correlation of individual contrasts between *conditions* (rhythmic-melodic *mTRF fits*) and the *PLV* for the “short-term”, but not the “long-term” model (Fig. 3B). This indicates the up-weighting of rhythmic predictions (and the down-weighting of pitch predictions) resulted in significant differences between the neural neural of these predictions in the “short-term” model, however, effects were weaker in the “long-term” model.

Additionally, GLMMs showed effects of *enjoyment*, with ratings being significantly positively related to the neural prediction tracking in both models (“short-term”: estimate=0.001, standard error=0.000, 95% CI=[0.000–0.001], pFDR<0.001) (“long-term”: estimate=0.000, standard error=0.000, 95% CI=[0.000– 0.001], pFDR<0.001) (S2-3). The same effect applies to *familiarity* ratings (“short-term”: estimate= 0.001, standard error=0.000, 95% CI=[0.001–0.001], pFDR<0.001) (“long-term”: estimate=0.001, standard error=0.000, 95% CI=[0.000–0.001], pFDR<0.001) (S2-3). The marginal and conditional R^2^ of the whispering GLMM were 0.098 and 0.332 for the “short-term” and 0.062 and 0.294 for the “long-term” model.

Similar effects were observed in the “tapping” GLMMs (see S7-9).

### Effect of musical sophistication on neural tracking of musical predictions

In the “whispering” GLMMs, no effects of the *MSI* were observed. We found a significant interaction effect of *MSI x condition* only in the “tapping” GLMM for “short-term” predictions (S8). The effect had the same direction as auditory-motor synchronization strength (i.e. higher musical sophistication relates to a relative up-weighting of rhythmic predictions). Furthermore, most models yielded a significant three-way interaction ((*MSI* x *PLV*) x *condition*) that illustrates how the slopes of the effect of *PLV* on *mTRF fit* is modified differently between conditions by the *MSI* (for detailed results e.g. S2, S8, S9). Yet, three-way interactions need to be interpreted cautiously, and the direction of the three-way interactions was not consistent across model and therefore we refrain from further interpretation.

## Discussion

The perception of music is facilitated by predictive processes which furthermore may be related to our tendency to move to and enjoy music. During naturalistic music listening, we provide evidence for differences in the temporal acoustic processing and the weighting of higher-level (short- and long-term) rhythmic and melodic predictions related to an individuals’ motor system recruitment strength. Individuals with strong auditory-motor synchronization behavior exhibited stronger neural tracking of lower-level temporal aspects of the music signal (the acoustic envelope and note onsets). This strongly suggests that auditory-motor networks are implicated in the processing of musical rhythms, also during a pure listening task that involves no motor responses. Interestingly, auditory-motor synchronization strength was also related to differential weighting of rhythmic and melodic predictions. High auditory-motor synchronizers neurally tracked rhythmic short-term and long-term predictions more strongly compared to melodic predictions, while low synchronizers showed no such preference. We further were able to show that this pattern is expressed more strongly for the use of short-term predictions (predictions based on local context only). Finally, the perceived familiarity with and enjoyment of the music predicted the neural tracking strength. Higher enjoyment was related to lower neural tracking of acoustic temporal features, while both high enjoyment and familiarity were associated with stronger tracking of melodic and rhythmic predictions. Our findings provide novel insights suggesting that individuals who can recruit the motor system more strongly, seem to prefer to neurally track temporal aspects of the music at different processing levels. In other words, this is expressed by advanced neural tracking of temporal aspects during lower-level acoustic processing and a selective up-weighting of rhythmic over melodic high-level predictions.

### Auditory-motor synchronization predicts neural tracking of temporal aspects of the music acoustics

We found significant neural tracking of temporal aspects of the music acoustics (acoustic envelope and note onset) across the whole brain. The neural tracking was strongest at fronto-central sites, likely indicating sources in the auditory and frontal cortex (e.g. 44–46). Our findings are in line with previous research that reported neural tracking of acoustic temporal features of music, including the acoustic envelope and note onsets (16, 23, 24, 78). Particularly, acoustic envelope tracking has been interpreted as reflecting temporal segmentation or rhythmic processing of speech and music (79). Crucially, we found higher neural tracking of acoustic temporal features in individuals with high-auditory motor synchronization strength, as hypothesized. In several studies, auditory-motor synchronization strength has been positively associated with improved temporal processing (31, 33, 34, 36, 38). Additionally, auditory-motor training was shown to improve speech perception (80). Our findings suggest that such faciliatory effects may occur at the level of the processing of acoustic temporal features during music listening. This is in line with a recent finding that neural envelope tracking during music listening is related to finger tapping beat synchronization performance (25). Note that we only found effects of auditory-motor synchronization in the “whispering” but not the “tapping” task. A difference between the finger tapping task in Keitel et al. (25) and our study is that the tapping task was relatively harder in the study by Keitel et al., possibly resulting in a better sensitivity to identify individual differences. In our study, the “whispering” compared to the “tapping” auditory-motor synchronization measure seemed more suitable to predict neural tracking of acoustic features.

### Auditory-motor synchronization predicts weighting of rhythmic and melodic predictions

Short-term and long-term rhythmic and melodic predictions were significantly neurally tracked at a fronto-central electrode cluster, suggesting that individuals extracted and utilized musical predictions. Importantly, both, melodic and rhythmic, predictions were tracked on top of the acoustic temporal tracking (i.e., added a significant gain in *mTRF fit* after controlling for acoustic features). Our findings are in line with previous studies showing the neural tracking of melodic predictions (23, 24, 81), and a study showing the neural tracking of rhythmic predictions (23). Our findings suggest that musical predictions were extracted based on both, local-context (“short-term” model) as well as on long-term priors (“long-term” model). This is in line with a previous study that showed the neural tracking of both short-term and long-term melodic predictions (23). In both models, rhythmic predictions were neurally tracked significantly stronger than melodic predictions. In contrast, the previous study (26) that investigated rhythmic and melodic prediction tracking found no overall difference between the neural tracking of these predictions. The discrepancy to our study likely is related to differences in the study sample and paradigm (e.g., [27] contained a smaller sample that included professional musicians, and stimulus repetitions). Note that we replicated the stronger tracking of rhythmic vs pitch predictions in our control analysis with the same IDyOM settings as used in [27] (see S11).

Importantly, the neural tracking of rhythmic and melodic predictions interacted with the auditory-motor synchronization strength (for both, “whispering” and “tapping” tasks). That is, with an increased auditory-motor synchronization strength, individuals exhibited an increased relative up-weighing of rhythmic predictions over melodic predictions. Albeit present for both, “short-term” and “long-term” prediction models, the effect was particularly pronounced in “short-term” prediction model. We showed for the first time -to the best of our knowledge-that individuals who more strongly spontaneously recruit the motor system, use it not only to improve the acoustic temporal processing of music, but also to more strongly utilize short-term rhythmic musical predictions as well as predictions based on long-term knowledge. At the same time, individuals with high auditory-motor synchronization strength down-weighted the neural tracking of melodic predictions. Thus, although high auditory synchronizers showed a relative up-weighting of rhythmic over pitch predictions their overall neural prediction tracking was not increased compared to low synchronizers. Bases on the previous literature (33, 34, 36, 38), we hypothesize that increased rhythm processing to be advantageous for music perception. Alternatively, the findings, however, may indicate weighting preferences - not related to processing abilities. Future research needs to further investigate this in an active paradigm where conclusions about which weighting is the most advantageous for music perception are possible.

### Effects of musical sophistication on acoustic temporal and musical prediction tracking

We included musical sophistication as control variable to ensure auditory-motor synchronization effects do not merely reflect enhanced musical sophistication. As expected from previous research (35), we found a positive correlation between musical sophistication and auditory-motor synchronization strength. However, musical sophistication was not predictive of acoustic feature tracking. While studies that compared musicians and non-musicians (16, 24) found a positive effect of musical training on acoustic envelope tracking, such correlation was present in one study that implemented musical sophistication as continuous variable amongst non-musicians (25), yet with no effect in another study (70).

Regarding the neural tracking of musical predictions, the evidence for the influence of musical sophistication was inconsistent in our study. We observed a significant effect in only some of the computational prediction models (see e.g. S8), suggesting a similar effect as auditory-motor synchronization strength (i.e. higher musical sophistication relates to a relative up-weighting of rhythmic predictions). A recent finding (32), furthermore, suggested a trade-off effect in favor of the neural tracking of pitch predictions over semantic predictions (in song) for individuals with high musical sophistication. As rhythmic prediction tracking was not explored, however, a direct comparison with our findings is not possible. In summary, our findings suggest that musical sophistication is a weaker predictor of neural tracking compared to auditory-motor synchronization (38). It seems to rather predict higher-level musical prediction tracking than acoustic tracking, albeit with the same direction of the effects as observed for auditory-motor synchronization.

### Familiarity, enjoyment and working memory effects on acoustic and long-term prediction tracking

Finally, the control variables of perceived familiarity and enjoyment predicted neural tracking strength. Higher enjoyment resulted in lowered neural tracking of acoustic temporal features, while both, enjoyment and familiarity, were associated with stronger neural tracking of musical predictions. Possibly, when individuals were not capable of neurally tracking higher-level predictions, they rather focused on lower-level acoustics instead, which, in turn, might have been related to lower enjoyment. Conversely, individuals who were able to track the predictions more closely, also enjoyed the music more as they could engage emotionally more strongly. However, in contrast, Keitel et al. (25) found the opposite effect of enjoyment on the neural tracking of acoustics, whereas Weineck et al. (70) found no effect. Taken together, this shows that further research is required to clarify the effect of enjoyment on neural tracking of acoustic features. While stronger acoustic tracking for more familiar music has been previously shown (70), we only found increased musical prediction tracking with higher familiarity, which has not been investigated previously. Generally, previous findings are heterogeneous and comparing ratings across studies can prove difficult as studies vary with respect to the sample and paradigm (e.g. stimulus repetitions, expert musicians etc.) and the rating scales (e.g. 69, 70, 80).

## Conclusion

Our findings provide novel insights suggesting that individuals who strongly recruit the motor system show advanced lower-level acoustic processing of temporal features (acoustic envelope and note onset tracking). Beyond that, we provided evidence for a relative up-weighting of rhythmic over melodic predictions in individuals with strong motor system recruitment. Crucially, we were able to show this effect for both, short-term predictions (based on local context only) and long-term predictions (based on long-term musical enculturation). Such focus on rhythmic processing is expected to be advantageous for music perception, however, future research is required to test such proposal in a task-related paradigm.

## Materials and Methods

The study protocol and analyses were preregistered on asPredicted.org (https://aspredicted.org/hqr8-48fv.pdf). Deviations from the preregistered procedure are discussed in the Supplement.

### Participants

The data of 45 participants were recorded. Five participants were excluded from the analyses for data quality reasons (lack of wakefulness, technical issues, missing data). Hence, the data set of 40 participants was analyzed (21 female, 19 male; the German “Geschlecht” was indicated according to self-report, M(age)=27.50y, SD(age)=4.04y, range(age)=21-35y). All participants reported to be neurologically healthy, without psychiatric disorders, dyslexia or dyscalculia, right-handed and to have normal, uncorrected hearing. None indicated to be a professional musician. All participants gave written informed consent and received monetary compensation for their participation. All experimental procedures were ethically approved by the Ethics Council of the Max Planck Society (No. 2017_12). Following the SSS-test protocol (67), another four data sets were excluded for all analyses that included the “whispering” *PLV* (N=36) due to incongruency across trials (within a 95% confidence interval). No incongruencies were found for the “tapping” task. For all analyses that included the *mTRF fits*, additional exclusions were executed based on outlier criteria (2.5 standard deviations from the mean), resulting in one exclusion for the acoustic tracking LMMs (N=39 for tapping GLMMs, N=35 for whispering GLMMs).

### General procedure

After the EEG cap was administered, the participants were seated in a sound attenuated booth in front of a computer monitor. The experiments were programmed in Matlab R2021a, using Psychophysics Toolbox extensions (83, 84). The stimulus presentation and response recordings were performed using a Windows PC. All auditory stimuli were presented binaurally with Ethymotic Research 3C in-ear headphones with E-A-RLINK foam tips attached. The study consisted of one session that lasted on average 140 min, including EEG preparation. The order of the tasks took place as follows: naturalistic music listening task (45 min, including 15 min for ratings after each musical piece and breaks), Digit Span Test (15 min), perception-production synchronization task (the order of the finger tapping and whispering synchronization task, SSS test, was randomized across participants) (30 min), and an online survey including questions about task specific, and demographic information as well as the Goldsmith Musical Sophistication Index (MSI, (68)) (Fig. 1A).

### Naturalistic music listening task

#### Stimuli

11 musical pieces were presented during the music listening task (total duration=28.19 min, details in S12). Each experimental block consisted of one musical piece. The total stimulus time meets the recommendation for a within-subject mTRF analysis of continuous auditory stimuli (85). All melodies were continuous, monophonic melody lines from existing musical pieces for violin, flute, or oboe and were synthesized using grand piano sounds (to avoid effects of different timbre) in MuseScore3 (Version 3.6, MuseScore BVBA), exported as MIDI-file for further analysis as well as WAV-file (sampled in 44.1 kHz) for stimulus presentation. The tempo was fixed at a set tempo for each piece (M=102.46bpm, SD=30.22bpm, range=47-144bpm).

#### Procedure

The participants were instructed to attentively listen to the music while looking at the fixation cross on the computer screen. The participants started each trial by pressing a key on the keyboard (sound started after 2 seconds). After each block, participants rated the familiarity (1: unknown to 7: very familiar) and the enjoyment (1: did not enjoy) to 7: enjoyed thoroughly)) of the previous block to ensure attentiveness to the pieces and as measures used as control variables for the GLMM analyses.

### EEG data acquisition and preprocessing

EEG data were recorded from 64 electrode positions using actiCAP (by BrainProducts GmbH, Gilching Germany). The signal was amplified and digitized at 1 kHz with BrainAmp DC (by BrainProducts GmbH, Gilching Germany). Additionally, 3 electrooculogram electrodes were placed in the participants’ faces to record eye movement with the BrainAmp ExG (by BrainProducts GmbH, Gilching Germany). BrainVisionRecorder (86) was used to record the EEG data on a Windows PC. For preprocessing of the EEG data, we used the Python based toolbox “mne”. Firstly, the data were re-referenced to the average of the two mastoid channels to emphasize the EEG response to auditory stimuli (87). Then, we filtered the signal digitally between 0.5 and 30 Hz. We further performed independent component analysis (ICA) to identify and remove eye movement alongside cardiac activity related artifacts, bad channels were interpolated (spherical spline interpolation, on average 1,9 per participant). Lastly, the data were epoched and downsampled to 128 Hz for subsequent analyses. All data were z-score normalized across channels for mTRF analyses (62).

### Quantifying musical predictions

Musical predictions have been previously quantified based on computational modeling of *Surprisal* and *Entropy* using variable-order Markov (24–27, 29) or Transformer models (24, 30) and combined them with neural tracking approaches, such as multivariate temporal response functions (31–33). To estimate temporal onset (rhythmic) and pitch (melodic) predictions of the musical pieces (*Surprisal* and *Entropy*), the Information Dynamics of Music (IDyOM, (88)) model was used. We computed two IDyOM models: the “short-term” model and the “long-term” model (23, 66). While the “short-term” model only uses the local context, the “long-term” model is pre-trained on a Western Classical Music Corpus (consisting of 903 melodies, for details see Supplements). As order-bound of the n-grams for each model an n of 16 was chosen as it was shown to represent listeners’ experience (23, 24, 89). The number of k-folds for the cross-validation of the model was set to 10. The target viewpoints were set to “cpitch” and “onset” in all models. In our primary model, the source viewpoints were set to “cpint” and “ioi” with the aim of comparability due to the use of simple intervals as reference for both pitch and onset. We further computed two additional models, one with “cpint”, “ioi-ratio” and “posinbar” as a more complex rhythm representation as well as one with “cpitch” and “ioi-ratio” as in Bianco et al., 2024 (26) to compare results (S11).

### Multivariate temporal response functions

To assess individual differences in using musical predictions, multivariate temporal response functions (mTRFs (62–64)) were computed which are based on regularized linear regression, using the mTRFpy toolbox (90). The chosen forward encoding model predicts the EEG response to a given stimulus (by leave-one-out cross-validation). The predicted and actual EEG response are compared yielding a correlation factor (Pearson’s r) or “*mTRF fit*”, which we use as index for neural tracking. The time-lag window to estimate neural responses was set from -50 ms to 400 ms (e.g. 16, 25). The regularization parameter (λ) was optimized across all participants for each model separately. Stimulus onset responses (500ms after trial start) were cut prior to analyses.

Three different models were used for the mTRF analyses. The “acoustic” model contained the acoustic envelope (z-score normalized), and the half-wave-rectified first derivative of the envelope representing the note onsets (91) (z-score normalized). The other four models additionally (on top of the acoustic features) contained musical prediction measures (*Surprisal* (i.e. *Information Content)* and *Entropy)* for either pitch (melodic prediction model) or temporal onsets (rhythmic prediction model) from either the IDyOM “short-term” or “long-term” model. The *mTRF fits* of all models were normalized with a shuffled null-distribution (mean of 1000 randomized permutations) (cf. 88). For the “acoustic” model, the order of the stimulus features was shuffled across trials (derangements between EEG and stimulus features only). For the musical prediction models, the order of the prediction values within a trial was shuffled while keeping the acoustic features constant, to maintain model dimensions while subtracting the *mTRF fit* gained by low level acoustic features and fitted noise.

### Perception-production synchronization task

The auditory-motor synchronization strength was measured behaviorally in two tasks, where participants either whispered to syllable sequences (SSS test, (33, 67)) or finger tapped to piano tone sequences (adapted SSS test; (36)) in synchrony. The stimuli from a previous study were used (36), which followed the protocol for the explicit SSS test (67). For the whispering task the syllabic rate accelerated from 4.3-4.7 Hz (in 0.1 Hz steps), a rate that is optimal for speech comprehension. For the finger tapping task, the rate of piano tones accelerated from 1.92-2.08 Hz (in 0.04 Hz steps), a rate that typically yielded the highest performances in finger tapping synchronization tasks (97, 98, 100). The stimuli were generated using “MBROLA”/”Praat” (syllables, with a male German diphone database (de2)) and “MIDIUtil” (piano tones). The syllables were randomly taken from a pool of 12 syllables (with no consecutive repetition) that consisted of a consonant followed by a vowel sound. The piano tones ranged from C3-B3. Both stimuli were sampled in 44.1 kHz. The order of the whispering and finger tapping task was randomized across participants. In the beginning of each task, participants adjusted the loudness to not hear themselves whispering or tapping (70-90 dB SPL). After a training, participants performed two task trials (50 s syllable sequence, 120 s piano tone sequence), where they synchronized their production to a random sequence of syllables or piano tones. The speech signal was recorded with a Shure MX418 Microflex directional gooseneck condenser microphone. The finger tapping was recorded with a tapping microphone (Schaller Oyster S/P).

The phase-locking value (*PLV*) between the filtered, resampled (128 Hz) envelopes of the auditory signal (cochlear envelope: 180 Hz and 7.246 kHz (101)) and the recorded motor output (102) was computed to quantify auditory-motor synchronization strength using Matlab R2024a. Filters were applied based on the rate of the stimulus, according to the SSS test protocol (65, https://zenodo.org/records/6142988). The *PLV*s were calculated for certain overlapping time windows (5 s for speech, 11 s for tapping; overlap: 2 s for speech, 4.5 s for tapping) and then averaged over both trials within each condition (whispering or tapping). The *PLV* can range from 0 (no synchronization) to 1 (maximal synchronization).

### Digit Span test

In an auditory Digit Span test (forward and backward) series of digits between 1-9 were presented. At the end of each sequence, participants were instructed to enter the numbers in the correct order on the keyboard. Each condition was terminated when the participant failed to correctly repeat both trials of the same sequence length. The digit span score was calculated by averaging the length of the last correctly entered sequence of both conditions.

### Statistical analysis

We performed linear mixed-effects models (LMMs) to quantify the effects of auditory-motor synchronization (*PLV*), musical sophistication (*MSI*), and their interactions on neural tracking (*mTRF fits*). In the musical prediction tracking model, we added the fixed effect *condition*, coding for the rhythmic and melodic predictions to investigate weighting effects. A random structure was included for *subjects, trials* and *EEG-electrodes* (*channel*). Due to multicollinearity between the “whispering” and “tapping” *PLV*s, separate LMMs were fit. In case of heteroscedasticity of the dependent variable, generalized linear mixed models (GLMMs) (family=beta) was computed (after transformation of the dependent variable *mTRF fit*, 0<*mTRF fit*<1). Statistical analyses were implemented using R (Version 4.3.3) running on RStudio (Version 2023.12.1). The LMM analyses relied on the package “lmerTest” (Version 3.1.3). GLMMs were computed with the package “glmmTMB” (Version 1.1.11). Heteroscedasticity was tested with the “performance” package (Version 0.13.0). Plots were created in RStudio with “ggplot2” (Version 3.5.1) and “sjPlot” (Version 2.8.16) or in JupyterLab (Version 4.0.11 running on Python 3.12.4) with the Python based packages “seaborn” (Version 0.13.2) and “matplotlib” (Version 3.8.4).

To avoid inflation of p-values due to multiple comparisons, False Discovery Rate (FDR) correction (103) was applied to p-values in LMMs using the p.adjust() function (method=“fdr”) in RStudio.

Statistical analyses were carried out for electrode clusters that yielded a significant *mTRF fit* which were identified using threshold-free cluster enhancement non-parametric t-test (72, 73). Specifically, we used the function mne.stats.permutation_cluster_1samp_test, which is implemented in the “mne” toolbox for Python (Version 1.8.0) (two-tailed, p-threshold < 0.001, 1024 permutations).

## Acknowledgements

We thank the Max Planck Institute for Empirical Aesthetics for funding this project (N.H, C.A., F.U. J.M.R.) as well as Dr. Klaus Frieler for valuable advice on statistical analyses and Ana Clemente for insightful information during music information retrieval.

## Data sharing options

The anonymized data and analysis code will be available upon publication.

