## Supplements for "Auditory-motor synchronization determines the use of predictions in music perception"

**Statistical results for “whispering” GLMMs**

S1: Detailed results of the “whispering” GLMM predicting acoustic feature tracking.

At the top, the table is displaying the detailed GLMM output. The plot below depicts the effect of *enjoyment* on acoustic feature tracking.

| | *predicted mTRF fit* | | | | |
|---|---|---|---|---|---|
| *Predictors* | *Estimates* | *SE* | *CI (95%)* | *z-values* | *p* |
| (Intercept) | 0.107 | 0.008 | 0.090 – 0.123 | 12.623 | **0.000** |
| *PLV whisper* | 0.018 | 0.006 | 0.007 – 0.029 | 3.173 | **0.004** |
| *MSI* | -0.009 | 0.005 | -0.020 – 0.001 | -1.785 | 0.130 |
| *digitspan* | -0.001 | 0.005 | -0.011 – 0.009 | -0.255 | 0.799 |
| *familiarity* | -0.000 | 0.000 | -0.001 – 0.001 | -0.945 | 0.402 |
| *enjoyment* | -0.002 | 0.000 | -0.003 – -0.001 | -4.396 | **0.000** |
| *PLV whisper × MSI* | -0.008 | 0.005 | -0.018 – 0.003 | -1.451 | 0.206 |
| **Random Effects** | | | | | |
| $\sigma^2$ | 0.00 | | | | |
| $\tau_{00\ trial}$ | 0.00 | | | | |
| $\tau_{00\ subject}$ | 0.00 | | | | |
| $\tau_{00\ channel}$ | 0.00 | | | | |
| ICC | 0.84 | | | | |
| $N_{trial}$ | 11 | | | | |
| $N_{sub}$ | 35 | | | | |
| $N_{channel}$ | 62 | | | | |
| Observations | 23870 | | | | |
| Marginal $R^2$ / Conditional $R^2$ | 0.059 / 0.852 | | | | |

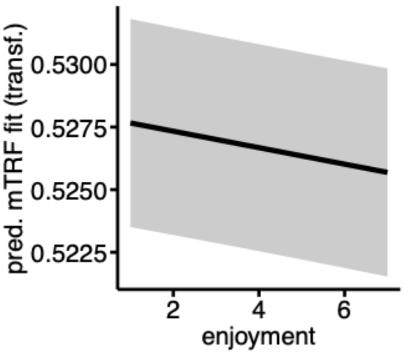

S2: Detailed results of the “whispering” GLMM predicting short-term prediction tracking.

At the top, the table is displaying the detailed GLMM output. The plots below depict the effect of *enjoyment* and *familiarity* on short-term prediction tracking. Below that, the plot shows the three-way interaction between *PLV*, *MSI* and *condition* on short-term prediction tracking.

| | ***predicted mTRF fit*** | | | | |
|---|---|---|---|---|---|
| *Predictors* | *Estimates* | *std. Error* | *CI (95%)* | *z-values* | *p(FDR)* |
| (Intercept) | 0.003 | 0.001 | 0.001 – 0.005 | 3.058 | **0.005** |
| *PLV whisper* | -0.002 | 0.001 | -0.003 – -0.000 | -2.063 | 0.061 |
| *MSI* | 0.000 | 0.001 | -0.001 – 0.002 | 0.423 | 0.739 |
| *condi [temporal]* | 0.006 | 0.000 | 0.006 – 0.007 | 26.518 | **0.000** |
| *digitspan* | 0.000 | 0.001 | -0.001 – 0.001 | 0.032 | 0.975 |
| *familiarity* | 0.001 | 0.000 | 0.001 – 0.001 | 8.531 | **0.000** |
| *enjoyment* | 0.001 | 0.000 | 0.000 – 0.001 | 4.960 | **0.000** |
| *PLV whisper × MSI* | 0.000 | 0.001 | -0.001 – 0.002 | 0.454 | 0.739 |
| *PLV whisper × condi [temporal]* | 0.002 | 0.000 | 0.002 – 0.003 | 10.423 | **0.000** |
| *MSI × condi [temporal]* | 0.000 | 0.000 | -0.000 – 0.001 | 0.707 | 0.659 |
| *(PLV whisper × MSI) × condi [temporal]* | -0.001 | 0.000 | -0.001 – -0.000 | -2.565 | **0.019** |
| **Random Effects** | | | | | |
| $\sigma^2$ | 0.00 | | | | |
| $\tau_{00\ trial}$ | 0.00 | | | | |
| $\tau_{00\ subject}$ | 0.00 | | | | |
| $\tau_{00\ channel}$ | 0.00 | | | | |
| ICC | 0.26 | | | | |
| $N_{trial}$ | 11 | | | | |
| $N_{sub}$ | 36 | | | | |
| $N_{channel}$ | 56 | | | | |
| Observations | 36432 | | | | |
| Marginal $R^2$ / Conditional $R^2$ | 0.098 / 0.332 | | | | |

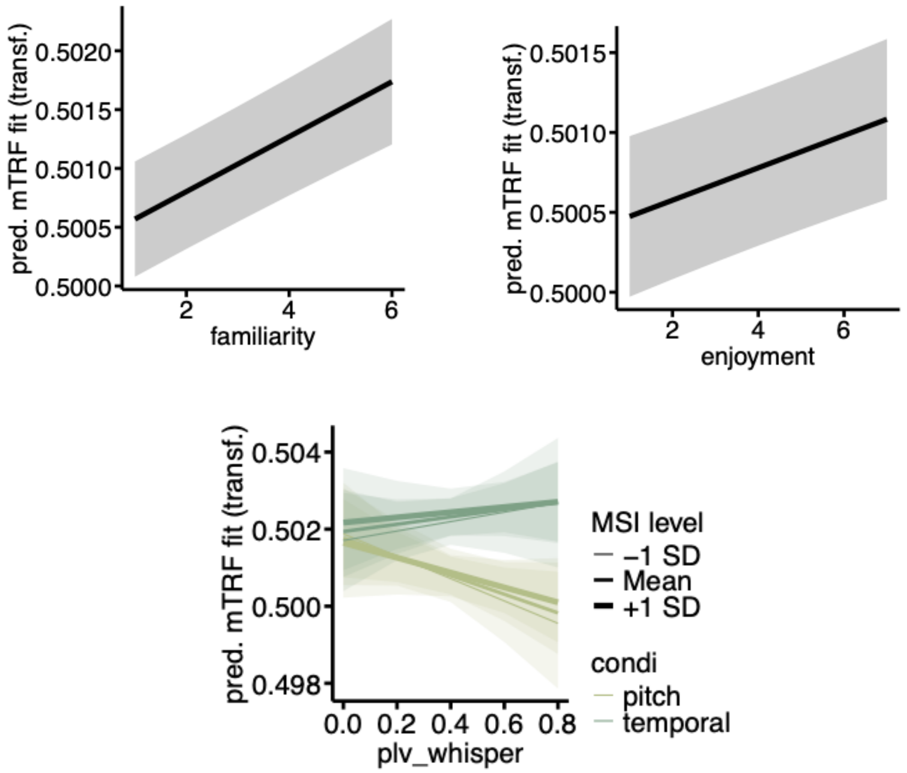

S3: Detailed results of the "whispering" GLMM predicting long-term prediction tracking.

At the top, the table is displaying the detailed GLMM output. The plots below depict the effect of *enjoyment* and *familiarity* on long-term prediction tracking.

| | ***predicted mTRF fit*** | | | | |
|---|---|---|---|---|---|
| *Predictors* | *Estimates* | *SE* | *CI (95%)* | *z-values* | p(FDR) |
| (Intercept) | 0.003 | 0.001 | 0.002 – 0.005 | 3.922 | **0.000** |
| *PLV whisper* | -0.001 | 0.001 | -0.003 – 0.000 | -1.871 | 0.084 |
| *MSI* | 0.001 | 0.001 | -0.001 – 0.002 | 0.797 | 0.520 |
| *condi [temporal]* | 0.003 | 0.000 | 0.003 – 0.004 | 17.416 | **0.000** |
| *digitspan* | 0.000 | 0.001 | -0.001 – 0.002 | 0.421 | 0.741 |
| *familiarity* | 0.001 | 0.000 | 0.000 – 0.001 | 6.089 | **0.000** |
| *enjoyment* | 0.000 | 0.000 | 0.000 – 0.001 | 3.799 | **0.000** |
| *PLV whisper × MSI* | 0.000 | 0.001 | -0.001 – 0.002 | 0.271 | 0.787 |
| *PLV whisper × condi [temporal]* | 0.001 | 0.000 | 0.001 – 0.001 | 4.816 | **0.000** |
| *MSI × condi [temporal]* | -0.000 | 0.000 | -0.001 – -0.000 | -1.983 | 0.074 |
| *(PLV whisper × MSI) × condi [temporal]* | 0.000 | 0.000 | 0.000 – 0.001 | 2.156 | 0.057 |

**Random Effects**

| | |
|---|---|
| $\sigma^2$ | 0.00 |
| $\tau_{00\ trial}$ | 0.00 |
| $\tau_{00\ subject}$ | 0.00 |
| $\tau_{00\ channel}$ | 0.00 |
| ICC | 0.25 |
| N trial | 11 |
| N sub | 36 |
| N channel | 45 |
| Observations | 31680 |
| Marginal $R^2$ / Conditional $R^2$ | 0.062 / 0.294 |

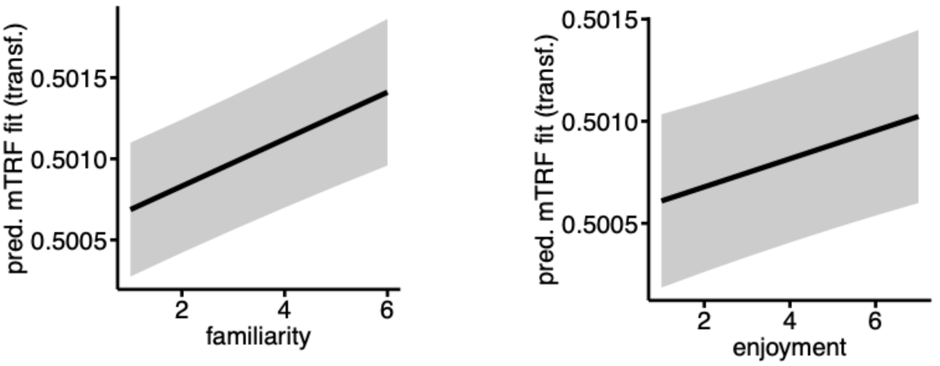

**Statistical results for “whispering“ GLMMs based on a pre-defined cluster of EEG-electrodes**

As a control analysis, we performed statistical analyses bof the mTRF fits at a pre-defined cluster of EEG-electrodes (e.g. 1–3). All other parameters of the analyses were kept the same like in our main analysis. The analyses yielded similar effects, as our anlyses with cluster-based regions of interest. The pre-defined cluster of EEG-electrodes consisted of fronto-central electrodes active during acoustic processing. Namely, Fz, Cz, FC3, FC1, C3, C1, FC2, FC4, C2, C4, F1, F2, FC5, C6, FC6, C5.

S4: Detailed results of the “whispering” LMM with channels in a pre-defined region of interest predicting acoustic feature tracking. Similar effects as in reported models using cluster-based electrode selection (statistical inference).
At the top, the table is displaying the detailed LMM output. The plots below depict the effect of *PLV* and *enjoyment* on acoustic feature tracking.

| | ***predicted mTRF fit*** | | | | |
|---|---|---|---|---|---|
| *Predictors* | *Estimates* | *SE* | *CI (95%)* | *z-values* | p(FDR) |
| *(Intercept)* | 0.058 | 0.136 | -0.209 – 0.325 | 0.427 | 0.743 |
| *PLV whisper* | 0.343 | 0.121 | 0.105 – 0.580 | 2.826 | **0.030** |
| *MSI* | -0.139 | 0.112 | -0.358 – 0.080 | -1.241 | 0.393 |
| *digitspan* | 0.039 | 0.109 | -0.176 – 0.253 | 0.353 | 0.743 |

| | | | | | |
|---|---|---|---|---|---|
| *familiarity* | -0.004 | 0.014 | -0.031 – 0.022 | -0.327 | 0.743 |
| *enjoyment* | -0.040 | 0.013 | -0.065 – -0.014 | -2.992 | **0.020** |
| *PLV whisper × MSI* | -0.143 | 0.112 | -0.363 – 0.077 | -1.272 | 0.393 |
| **Random Effects** | | | | | |
| $\sigma^2$ | 0.54 | | | | |
| $\tau_{00}$ subject | 0.36 | | | | |
| $\tau_{00}$ channel | 0.08 | | | | |
| $\tau_{00}$ trial | 0.00 | | | | |
| ICC | 0.46 | | | | |
| N trial | 11 | | | | |
| N sub | 34 | | | | |
| N channel | 16 | | | | |
| Observations | 5984 | | | | |
| Marginal $R^2$ / Conditional $R^2$ | 0.090 / 0.504 | | | | |

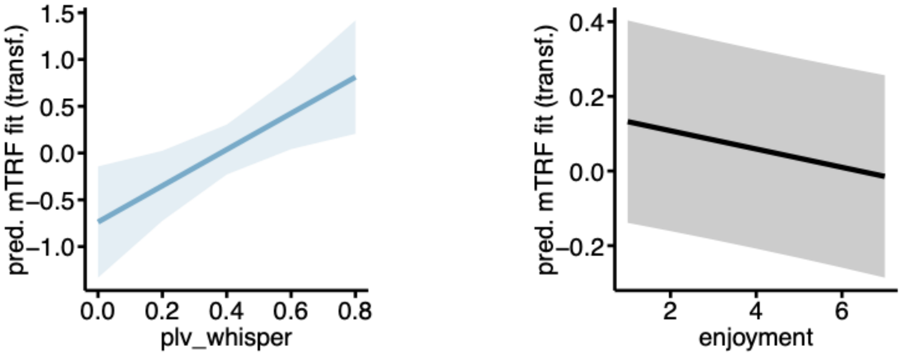

S5: Detailed results of the "whispering" GLMM with channels in a pre-defined region of interest predicting short-term prediction tracking. Similar effects as in reported models using cluster-based electrode selection (statistical inference).

At the top, the table is displaying the detailed GLMM output. The plot below on the left depicts the effect of *condition* on short-term prediction. The plot on the right-hand side shows the interaction between *condition* and *PLV* on short-term prediction tracking. The plots below show the effects of *familiarity* and *enjoyment* on short-term prediction tracking.

| | ***predicted mTRF fit*** | | | | |
|---|---|---|---|---|---|
| *Predictors* | *Estimates* | *std. Error* | *CI (95%)* | *z-values* | *p(FDR)* |
| (Intercept) | 0.005 | 0.001 | 0.003 – 0.008 | 4.734 | **0.000** |
| plv whisper | -0.002 | 0.001 | -0.004 – 0.000 | -1.958 | 0.092 |
| MSI | 0.001 | 0.001 | -0.001 – 0.002 | 0.840 | 0.551 |
| condi [temporal] | 0.007 | 0.000 | 0.007 – 0.008 | 22.466 | **0.000** |
| digitspan | 0.000 | 0.001 | -0.001 – 0.002 | 0.377 | 0.777 |

| | | | | | |
|---|---|---|---|---|---|
| familiarity | 0.001 | 0.000 | 0.001 – 0.001 | 4.983 | **0.000** |
| enjoyment | 0.001 | 0.000 | 0.000 – 0.001 | 3.327 | **0.002** |
| plv whisper × MSI | 0.000 | 0.001 | -0.002 – 0.002 | 0.223 | 0.823 |
| plv whisper × condi [temporal] | 0.002 | 0.000 | 0.002 – 0.003 | 7.078 | **0.000** |
| MSI × condi [temporal] | 0.000 | 0.000 | -0.000 – 0.001 | 0.570 | 0.695 |
| (plv whisper × MSI) × condi [temporal] | -0.001 | 0.000 | -0.001 – 0.000 | -1.674 | 0.148 |
| **Random Effects** | | | | | |
| $\sigma^2$ | 0.00 | | | | |
| $\tau_{00\ trial}$ | 0.00 | | | | |
| $\tau_{00\ subject}$ | 0.00 | | | | |
| $\tau_{00\ channel}$ | 0.00 | | | | |
| ICC | 0.30 | | | | |
| $N_{trial}$ | 11 | | | | |
| $N_{sub}$ | 35 | | | | |
| $N_{channel}$ | 16 | | | | |
| Observations | 12320 | | | | |
| Marginal $R^2$ / Conditional $R^2$ | 0.147 / 0.405 | | | | |

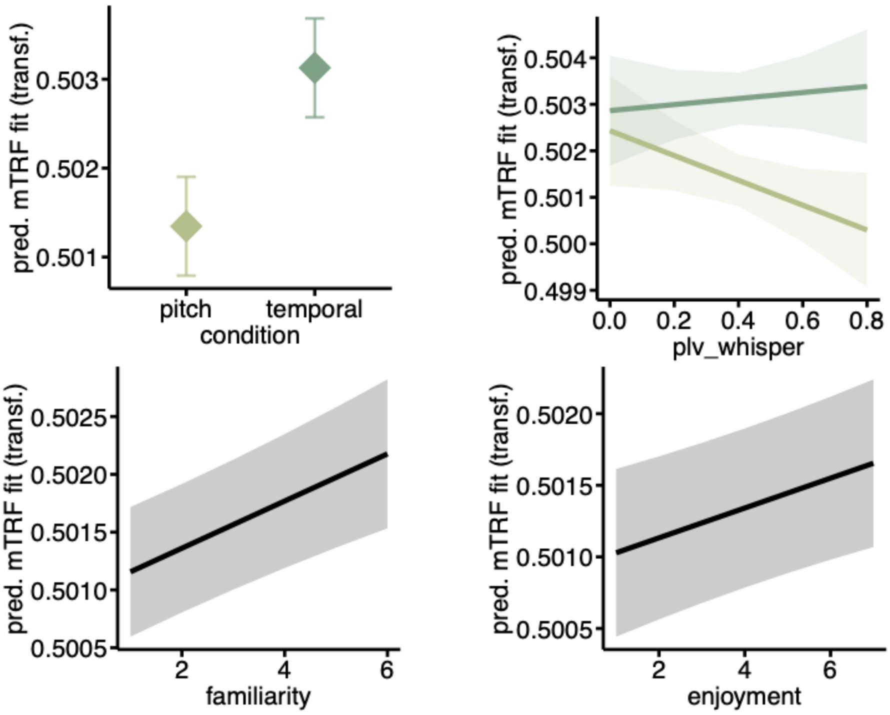

S6: Detailed results of the "whispering" GLMM with channels in a pre-defined region of interest predicting long-term prediction tracking. Similar effects as in reported models using cluster-based electrode selection (statistical inference).

At the top, the table is displaying the detailed GLMM output. The plot below on the left depicts the effect of *condition* on long-term prediction. The plot on the right-hand side shows the interaction between *condition* and *PLV* on short-term prediction tracking. The plots below depict the effects of *familiarity* and *enjoyment* on long-term prediction tracking.

| | ***predicted mTRF fit*** | | | | |
|---|---|---|---|---|---|
| *Predictors* | *Estimates* | *std. Error* | *CI (95%)* | *z-values* | *p(FDR)* |
| (Intercept) | 0.004 | 0.001 | 0.002 – 0.006 | 4.533 | **0.000** |
| *PLV whisper* | -0.001 | 0.001 | -0.003 – 0.000 | -1.814 | 0.128 |
| *MSI* | 0.001 | 0.001 | -0.001 – 0.002 | 0.961 | 0.463 |
| *condi [temporal]* | 0.004 | 0.000 | 0.004 – 0.005 | 16.269 | **0.000** |
| *digitspan* | 0.000 | 0.001 | -0.001 – 0.002 | 0.518 | 0.665 |
| *familiarity* | 0.001 | 0.000 | 0.000 – 0.001 | 3.032 | **0.007** |
| *enjoyment* | 0.000 | 0.000 | 0.000 – 0.001 | 2.578 | **0.022** |
| *PLV whisper × MSI* | 0.000 | 0.001 | -0.001 – 0.002 | 0.347 | 0.728 |
| *PLV whisper × condi [temporal]* | 0.001 | 0.000 | 0.000 – 0.001 | 3.200 | **0.005** |
| *MSI × condi [temporal]* | -0.000 | 0.000 | -0.001 – 0.000 | -1.459 | 0.227 |
| *(PLV whisper × MSI) × condi [temporal]* | 0.000 | 0.000 | -0.000 – 0.001 | 0.759 | 0.547 |
| **Random Effects** | | | | | |
| $\sigma^2$ | 0.00 | | | | |
| $\tau_{00\ trial}$ | 0.00 | | | | |
| $\tau_{00\ subject}$ | 0.00 | | | | |
| $\tau_{00\ channel}$ | 0.00 | | | | |
| ICC | 0.29 | | | | |
| $N_{trial}$ | 11 | | | | |
| $N_{sub}$ | 35 | | | | |
| $N_{channel}$ | 16 | | | | |
| Observations | 12320 | | | | |
| Marginal $R^2$ / Conditional $R^2$ | 0.092 / 0.354 | | | | |

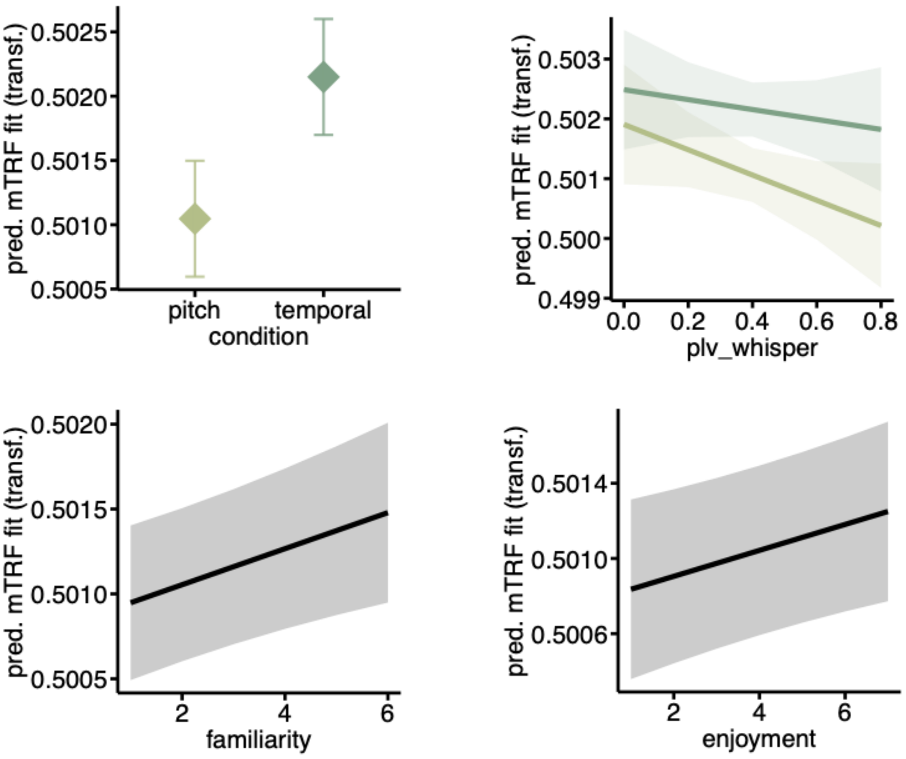

**<u>Statistical results for “tapping” GLMMs</u>**

S7: Detailed results of the “tapping” GLMM predicting acoustic feature tracking.
At the top, the table is displaying the detailed GLMM output. The plot below depicts the effect of *enjoyment* on acoustic feature tracking.

| | ***predicted mTRF fit*** | | | | |
|---|---|---|---|---|---|
| *Predictors* | *Estimates* | *SE* | *CI (95%)* | *z-values* | *p(FDR)* |
| (Intercept) | 0.103 | 0.009 | 0.086 – 0.120 | 11.791 | **0.000** |
| *PLV tap* | 0.008 | 0.007 | -0.006 – 0.023 | 1.144 | 0.509 |
| *MSI* | -0.005 | 0.006 | -0.016 – 0.006 | -0.909 | 0.509 |
| *digitspan* | 0.002 | 0.005 | -0.009 – 0.013 | 0.382 | 0.702 |
| *familiarity* | -0.001 | 0.001 | -0.002 – 0.001 | -0.973 | 0.509 |
| *enjoyment* | -0.002 | 0.000 | -0.003 – -0.001 | -4.370 | **0.000** |
| *PLV tap × MSI* | 0.002 | 0.005 | -0.007 – 0.011 | 0.417 | 0.702 |
| **Random Effects** | | | | | |
| $\sigma^2$ | 0.00 | | | | |
| $\tau_{00\ trial}$ | 0.00 | | | | |
| $\tau_{00\ subject}$ | 0.00 | | | | |
| $\tau_{00\ channel}$ | 0.00 | | | | |

| | |
|---|---|
| ICC | 0.85 |
| N trial | 11 |
| N sub | 35 |
| N channel | 62 |
| Observations | 23870 |
| Marginal $R^2$ / Conditional $R^2$ | 0.013 / 0.853 |

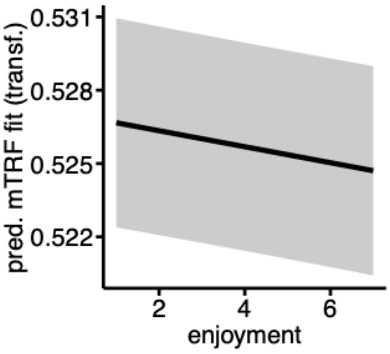

S8: Detailed results of the "*tapping*" GLMM predicting short-term prediction tracking.

At the top, the table is displaying the detailed GLMM output. The plots below depict the effect of *condition, familiarity* and *enjoyment* on short-term prediction tracking. The first two plots from the right-hand side in the bottom row show the interactions between *condition* and *PLV* as well as *MSI*, respectively, on short-term prediction tracking. The last plot in the bottom row depicts the three-way interaction between *PLV, MSI* and *condition* on short-term prediction tracking.

| | | | ***predicted mTRF fit*** | | |
|---|---|---|---|---|---|
| *Predictors* | *Estimates* | *std. Error* | *CI (95%)* | *z-values* | *p(FDR)* |
| (Intercept) | 0.003 | 0.001 | 0.001 – 0.005 | 3.477 | **0.001** |
| *PLV tap* | -0.000 | 0.001 | -0.002 – 0.002 | -0.114 | 0.963 |
| *MSI* | -0.001 | 0.001 | -0.002 – 0.001 | -0.723 | 0.574 |
| *condi [temporal]* | 0.006 | 0.000 | 0.005 – 0.006 | 25.583 | **0.000** |
| *digitspan* | 0.000 | 0.001 | -0.001 – 0.001 | 0.046 | 0.963 |
| *familiarity* | 0.001 | 0.000 | 0.001 – 0.001 | 8.542 | **0.000** |
| *enjoyment* | 0.001 | 0.000 | 0.000 – 0.001 | 4.940 | **0.000** |
| *PLV tap × MSI* | -0.000 | 0.001 | -0.002 – 0.001 | -0.831 | 0.558 |
| *PLV tap × condi [temporal]* | 0.001 | 0.000 | 0.000 – 0.001 | 3.515 | **0.001** |
| *MSI × condi [temporal]* | 0.001 | 0.000 | 0.001 – 0.001 | 4.734 | **0.000** |
| *(PLV tap × MSI) × condi [temporal]* | 0.001 | 0.000 | 0.000 – 0.001 | 3.382 | **0.001** |

| **Random Effects** | |
|---|---|
| $\sigma^2$ | 0.00 |
| $\tau_{00\ trial}$ | 0.00 |
| $\tau_{00\ subject}$ | 0.00 |
| $\tau_{00\ channel}$ | 0.00 |
| ICC | 0.26 |
| $N_{trial}$ | 11 |
| $N_{sub}$ | 36 |
| $N_{channel}$ | 56 |
| Observations | 36432 |
| Marginal $R^2$ / Conditional $R^2$ | 0.093 / 0.326 |

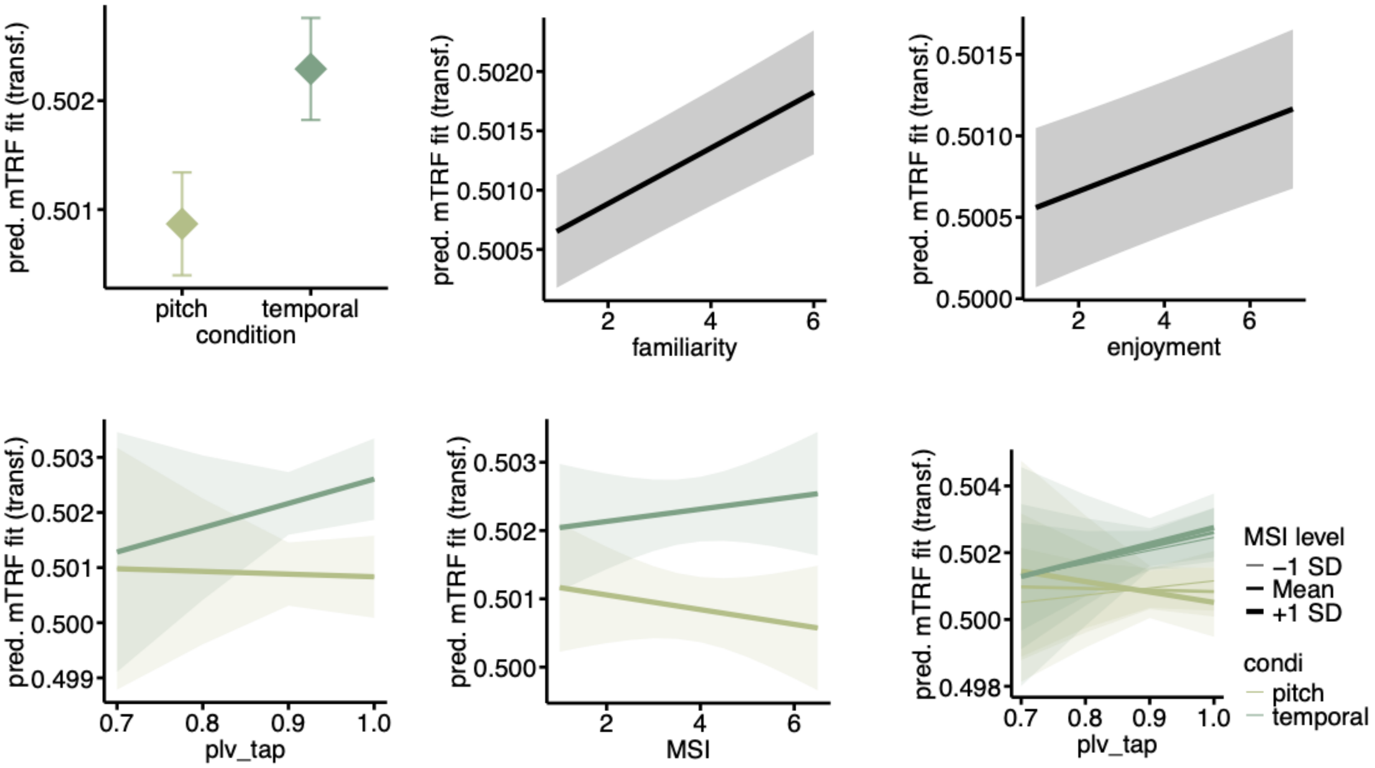

S9: Detailed results of the "tapping" GLMM predicting long-term prediction tracking**.**

At the top, the table is displaying the detailed GLMM output. The first plot from the right-hand side below depicts the effect of *condition* on long-term prediction. The second plot shows the interaction between *PLV* and *condition* on long-term prediction tracking. The last plot in the bottom row depicts the three-way interaction between *PLV, MSI* and *condition* on long-term prediction tracking.

| | ***predicted mTRF fit*** | | | | |
|---|---|---|---|---|---|
| *Predictors* | *Estimates* | *SE* | *CI (95%)* | *z-values* | p(FDR) |
| (Intercept) | 0.004 | 0.001 | 0.002 – 0.005 | 4.508 | **0.000** |
| *PLV tap* | -0.001 | 0.001 | -0.003 – 0.001 | -1.127 | 0.357 |

| | | | | | |
|---|---|---|---|---|---|
| *MSI* | -0.000 | 0.001 | -0.001 – 0.001 | -0.101 | 0.940 |
| *condi [temporal]* | 0.003 | 0.000 | 0.003 – 0.004 | 18.234 | **0.000** |
| *digitspan* | 0.000 | 0.001 | -0.001 – 0.001 | 0.410 | 0.833 |
| *familiarity* | 0.001 | 0.000 | 0.001 – 0.001 | 6.121 | **0.000** |
| *enjoyment* | 0.000 | 0.000 | 0.000 – 0.001 | 3.795 | **0.000** |
| *PLV tap × MSI* | -0.001 | 0.001 | -0.002 – 0.000 | -1.564 | 0.185 |
| *PLV tap × condi [temporal]* | 0.001 | 0.000 | 0.000 – 0.001 | 3.308 | **0.002** |
| *MSI × condi [temporal]* | -0.000 | 0.000 | -0.000 – 0.000 | -0.075 | 0.940 |
| *(PLV tap × MSI) × condi [temporal]* | 0.000 | 0.000 | 0.000 – 0.001 | 2.621 | **0.016** |
| **Random Effects** | | | | | |
| $\sigma^2$ | 0.00 | | | | |
| $\tau_{00}$ trial | 0.00 | | | | |
| $\tau_{00}$ subject | 0.00 | | | | |
| $\tau_{00}$ channel | 0.00 | | | | |
| ICC | 0.25 | | | | |
| N trial | 11 | | | | |
| N sub | 36 | | | | |
| N channel | 45 | | | | |
| Observations | 31680 | | | | |
| Marginal $R^2$ / Conditional $R^2$ | 0.057 / 0.290 | | | | |

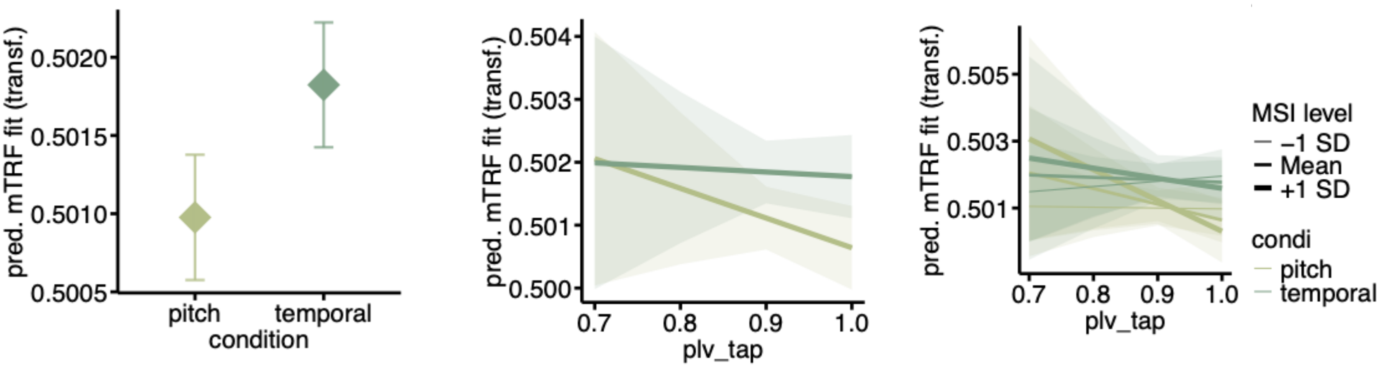

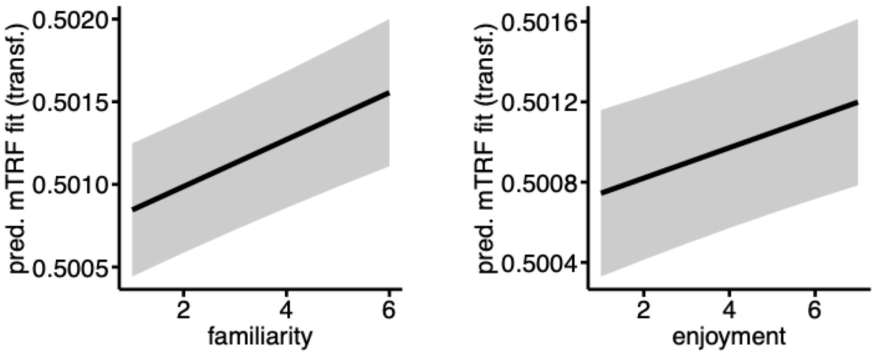

**S10: Distribution of HIGH and LOW audirory-motor synchronizer using k-means splitting.**
Histogram shows the number of participants in each group, amounting to 17 HIGHs and 17 LOWs. Visual inspection suggests the distribution varies from unimodality.

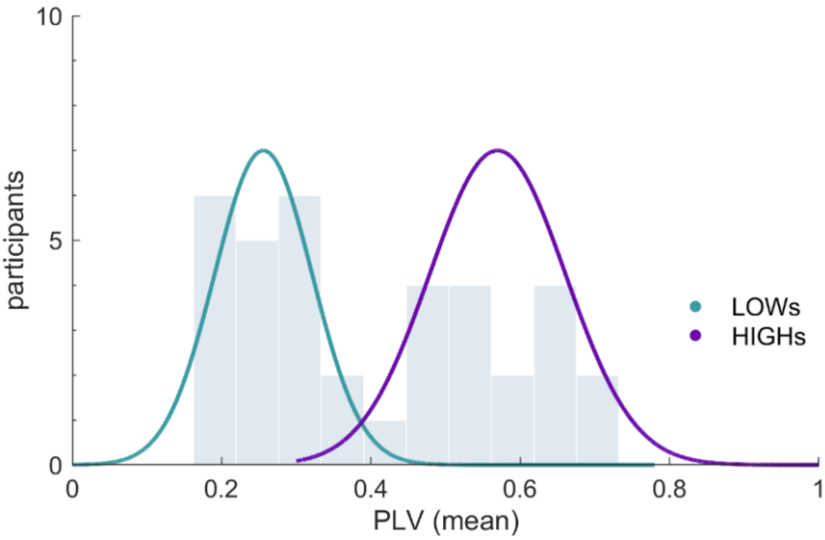

**Descriptive statistics of the control variables**

The ratings showed that the participants were not very familiar with the musical pieces (familiarity: M=1.95, SD=0.83, min=1.00, max=4.55) and that participants enjoyed them moderately (enjoyment: M=4.00, SD=1.02, min=1.73, max=6.00).

The mean auditory working memory capacity of participants was 5.69 digits (SD=0.76, min=4.5, max=7.5) measured as digit span. Participants showed a moderately high musical sophistication (MSI: M=3.78, SD=1.23, min=1.11, max=6.06) in our sample, which was expected as we did not recruit professional musicians.

**S11: Overview of statistical results from GLMMs based on different source-viewpoints for IDyOM.**

“cpint ioi-ratio posinbar” is a model that computes predictions based on more strongly musically informed temporal and pitch patterns, e.g. for rhythm the metrical structure is considered.“cpitch ioi-ratio” is the model used in a primary reference paper and was computed to compare results.
The table below depicts the estimates, and FDR-corrected p-values in brackets.
cpint: pitch interval
cpitch: pitch (on a chromatic scale)
ioi-ratio: inter-onset-interval (ioi) ratio (ioi divided by previous ioi)
posinbar: time offset from beginning of bar

| | cpint ioi-ratio posinbar | | | cpitch ioi-ratio | |
|---|---|---|---|---|---|
| Predictor | short-term | long-term | | short-term | long-term |
| *PLV whisper* | -0.002 (0.100) | -0.001 (0.186) | | -0.000 (0.861) | -0.000 (0.911) |
| *MSI* | 0.000 (0.806) | 0.000 (0.735) | | 0.000 (0.978) | 0.001 (0.911) |
| *condi [temporal]* | 0.007 (**0.000**) | 0.006 (**0.000**) | | 0.010 (**0.000**) | 0.010 (**0.000**) |
| *digitspan* | 0.000 (0.921) | -0.000 (0.949) | | 0.000 (0.978) | 0.000 (0.911) |
| *familiarity* | 0.000 (0.100) | -0.001 (**0.000**) | | 0.000 (0.754) | 0.000 (0.911) |
| *enjoyment* | 0.001 (**0.000**) | 0.002 (**0.000**) | | 0.001 (**0.000**) | 0.000 (0.057) |
| *PLV whisper × MSI* | 0.000 (0.806) | 0.000 (0.949) | | 0.001 (0.586) | -0.001 (0.911) |
| *PLV whisper × condi [temporal]* | 0.002 (**0.000**) | 0.001 (**0.017**) | | 0.001 (**0.036**) | -0.000 (0.911) |
| *MSI × condi [temporal]* | 0.000 (0.232) | 0.000 (0.353) | | 0.001 (**0.067**) | -0.001 (0.911) |
| *(PLV whisper × MSI) × condi [temporal]* | -0.000 (0.386) | -0.001 (**0.004**) | | -0.000 (0.107) | 0.001 (0.537) |

**S12: Details on musical stimuli**, all pieces were presented with synthesized Grand Piano sound.

| Composer | Compositions | Year | Key | Time Signature | Tempo (bpm) | Duration (s) | Notes |
|---|---|---|---|---|---|---|---|
| Antonio Vivaldi | Oboe Sonata in C minor, RV 53, 2nd movement | ca. 1703-1740 | C minor | 4/4 | 80 | 162.468 | 618 |
| Johann Sebastian Bach | Violin Sonata in B minor, BWV 1014, 2nd movement | 1717-23 | B minor | 4/4 | 120 | 192.936 | 506 |
| Johann Sebastian Bach | Partita in A minor, BWV 1013, 4th movement: Bourrée anglaise | 1722–23 | A minor | 2/4 | 80 | 133.799 | 529 |

| | | | | | | | |
|---|---|---|---|---|---|---|---|
| Friedrich Kuhlau | 3 Fantasies for Solo Flute, Op.38, Fantasy No. 3, 2nd movement | 1821 | C major | 4/4 | 116 | 136.008 | 499 |
| Friedrich Kuhlau | 3 Fantasies for Solo Flute, Op.38, Fantasy No. 1, 2nd movement | 1821 | D minor | 4/4 | 144 | 137.478 | 571 |
| Friedrich Kuhlau | 3 Fantasies for Solo Flute, Op.38, Fantasy No. 2, 2nd movement | 1821 | G minor | 3/4 | 120 | 185.211 | 809 |
| Franz Schubert | Arpeggione Sonata, D.821, 1st movement | 1824 | A minor | 4/4 | 110 | 133.608 | 526 |
| Johann Sebastian Bach | Violin Partita No.3 in E major, BWV 1006, 3rd movement: Gavotte en Rondeau | 1720 | E major | 4/4 | 140 | 176.528 | 642 |
| Johann Sebastian Bach | Violin Partita No.2 in D minor, BWV 1004, 1st movement: Allemande | 1720 | D minor | 4/4 | 47 | 163.731 | 540 |
| Johann Sebastian Bach | Partita in A minor, BWV 1013, 3rd movement: Sarabande | 1722–23 | A minor | 3/4 | 70 | 118.155 | 301 |
| Johann Sebastian Bach | Partita in A minor, BWV 1013, 2nd movement: Corrente | 1722–23 | A minor | 4/4 | 100 | 151.469 | 891 |
| | | | | | **total** | 1961.391 | 6432 |
| | | | | | **M** | 153s | 584.73 |
| | | | | | **SD** | 24.24s | 158.52 |
| | | | | | **range** | 118.16-192.94s | 301-891 |

Musical pieces from different composers and musical eras were chosen to allow for stylistic variety. The long-term model of IDyOM was trained with a corpus of Western Classical music. It included a total of 738 melodies: 566 German folk melodies (dataset “fink” from the Essen Folk Song Collection (ESFC) by Schaffrath (4), https://kern.humdrum.org/cgi-bin/browse?l=essen/europa/deutschl/fink), 152 folk songs and ballads from Nova Scotia, Canada (collecte 1928-1932 by Creighton (5), https://kern.ccarh.org/cgi-bin/ksbrowse?s=nova), and 185 chorales by J. S. Bach (BWV 253 to BWV 438, https://kern.ccarh.org/cgi-bin/ksbrowse?type=collection&l=/musedata/bach/chorales).

**Deviations from the preregistration**

This study was preregistered under https://aspredicted.org/hqr8-48fv.pdf. In the process of analyses, the following adjustments were made.

Deviations in the Methods: mTRF model baseline

Instead of subtracting the mTRF for each channel of the “acoustic” model from the prediction models, we used a more sophisticated approach where we used a shuffled null-distribution as baseline. This way, we could account for dimensionality of the mTRF models. In other words, this ensures that *mTRF* *fit* differences are attributable to the model design rather than mismatched dimensions of the models as the amount of data alone can increase the likelihood of fitting noise. This approach, furthermore, allowed to baseline clean the “acoustic” model itself and perform follow-up statistics on the acoustic model. Please find the details regarding baseline cleaning in the Methods.

Deviations in the Analyses: correlation of “whispering” and “tapping” *PLV*s

As opposed the expectation in the pre-registration, *PLVs were* significantly correlated. As this would have caused multicollinearity issues if both were included within one GLMM, separate GLMMs were computed.

Deviations in the Analyses: PLV groups as predictor

In the preregistration it was planned to additionally to the GLMMs that we report in the manuscript, as a post-hoc analysis fit GLMMs to predict the contrast between rhythmic and melodic *mTRF fits* with the *PLV as categorical fixed predictor* (c *PLV groups,* HIGHs/LOWs) . This analysis may have less power than the analysis with *PLV* included as continuous variable. Furthermore, it wdoes not make it possible to perform a similar approach with the tapping data, as these show no bimodal distribution. The analysis with *PLV groups* shows similar effects, reflected in fixed effect of *sss group* which remains as trend after FDR correction.

Illustrative example of “whispering“ GLMM for “short-term“ predictions. Detailed results. *sss group [1]* refers to the so-called HIGHs. The positive fixed effect suggests an up-weighting of rhythmic over melodic predictions on a group level.

| | ***predicted mTRF fit contrast (temporal-pitch)*** | | | | | |
|---|---|---|---|---|---|---|
| *Predictors* | *Estimates* | *SE* | *CI (95%)* | *z-values* | *p* | *p(FDR)* |
| (Intercept) | 0.004 | 0.001 | 0.002 – 0.007 | 3.138 | **0.002** | 0.006 . |
| *sss group [1]* | 0.004 | 0.002 | 0.000 – 0.007 | 2.080 | **0.037** | 0.066 . |
| *MSI* | 0.001 | 0.001 | -0.001 – 0.004 | 0.924 | 0.355 | 0.498 |
| *digitspan* | 0.000 | 0.001 | -0.001 – 0.002 | 0.282 | 0.778 | 0.778 |
| *enjoyment* | 0.001 | 0.000 | 0.000 – 0.001 | 2.266 | **0.023** | 0.055 . |
| *familiarity* | 0.001 | 0.000 | 0.001 – 0.002 | 4.509 | **<0.001** | **0.000** |
| *sss group [1] × MSI* | -0.001 | 0.002 | -0.004 – 0.003 | -0.501 | 0.616 | 0.719 |
| **Random Effects** | | | | | | |
| $\sigma^2$ | 0.00 | | | | | |
| $\tau_{00\ trial}$ | 0.00 | | | | | |
| $\tau_{00\ subject}$ | 0.00 | | | | | |

| | |
|---|---|
| $\tau_{00\ channel}$ | 0.00 |
| ICC | 0.16 |
| $N_{trial}$ | 11 |
| $N_{sub}$ | 36 |
| $N_{channel}$ | 36 |
| Observations | 14256 |
| Marginal $R^2$ / Conditional $R^2$ | 0.045 / 0.202 |

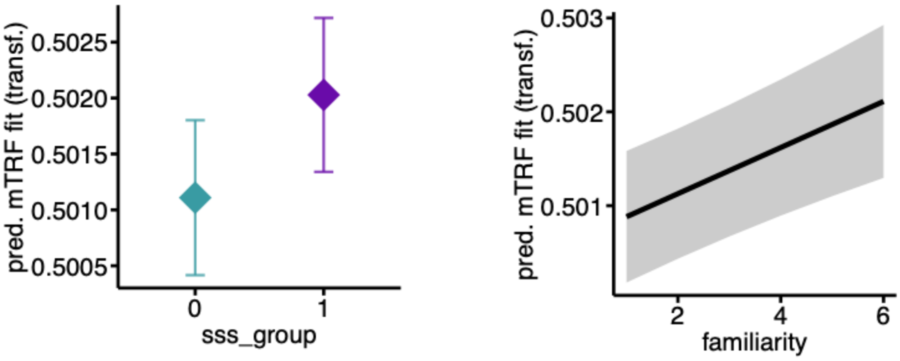
